# Sleep problems in preschoolers with autism spectrum disorders are associated with sensory sensitivities and thalamocortical overconnectivity

**DOI:** 10.1101/2021.01.15.426899

**Authors:** A.C. Linke, B. Chen, L. Olson, C. Ibarra, C. Fong, S. Reynolds, M. Apostol, M.K. Kinnear, R.-A. Müller, I. Fishman

## Abstract

Projections between the thalamus and sensory cortices are established early in development and play an important role in sleep regulation as well as in relaying sensory information to cortex. Atypical thalamocortical functional connectivity frequently observed in children with autism spectrum disorders (ASD) might therefore be linked to sensory and sleep problems common in ASD. Here we investigated the relationship between auditory-thalamic functional connectivity measured during natural sleep fMRI, sleep problems, and sound sensitivities in 70 toddlers and preschoolers (1.5 to 5-year-olds) with ASD compared to a matched group of 46 typically developing (TD) children. In children with ASD, sleep problems and sensory sensitivities were positively correlated, and increased sleep latency was associated with overconnectivity between the thalamus and auditory cortex in a subsample with high quality MRI data (n=29). Additionally, auditory cortex BOLD signal amplitude was elevated in children with ASD, potentially reflecting reduced sensory gating or a lack of auditory habituation during natural sleep. These findings indicate that atypical thalamocortical functional connectivity can be detected early in development and may play a crucial role in sleep problems and sensory sensitivities in ASD.

## 1. Introduction

Autism spectrum disorders (ASD) are neurodevelopmental disorders characterized by socio-communicative deficits, restricted and repetitive behaviors and interests, and sensory sensitivities (American Psychiatric Association, 2013). There is no known single etiology for ASD, but increasing evidence suggests that autism originates early in development, likely in utero (Courchesne et al., 2020; Gazestani et al., 2019; Grove et al., 2019). While a mechanism has not been established, atypical structural and functional brain organization is commonly observed in ASD (Holiga et al., 2019; Hull et al., 2017; Müller, 2014). The formation and maturation of connections between the thalamus and cerebral cortex (i.e., thalamocortical connections), mediated by the subplate, plays a crucial part in early cortical development (Constantinople and Bruno, 2013; Kanold and Luhmann, 2010; Kostović et al., 2011; Kostovic and Judas, 2010; O’Leary and Nakagawa, 2002). This process peaks during the third gestational trimester and – when disrupted – can result in distorted topographic cortical organization with cascading consequences for postnatal sensory processing, motor learning, and cognitive development (for reviews see Hadders-Algra, 2018; Hoerder-Suabedissen and Molnár, 2015; Kanold, 2019; Luhmann et al., 2018). Genetic predisposition (Bai et al., 2019; Colvert et al., 2015; Sandin et al., 2017) and to a lesser extent environmental factors such as maternal infection during pregnancy (al-Haddad et al., 2019; Lombardo et al., 2018), prematurity, or other pregnancy or birth complications (Agrawal et al., 2018; Chien et al., 2019; Getahun et al., 2017) constitute known risk factors for ASD, and have been shown to interfere with subplate neuron function and development of thalamocortical projections (Caubit et al., 2016; Hoerder-Suabedissen et al., 2013; Hoerder-Suabedissen and Molnár, 2015; Kanold, 2003; Materna et al., 2008; McClendon et al., 2017; McQuillen et al., 2003; Mikhailova et al., 2017; Toulmin et al., 2015). Abnormal function of subplate neurons during prenatal brain development has also been detected in autism (Hoerder-Suabedissen et al., 2013; Hutsler and Avino, 2015; Hutsler and Casanova, 2015; Kanold, 2009; Luhmann et al., 2018; Nagode et al., 2017; Serati et al., 2019; Wess et al., 2017), and focal laminar disorganization hypothesized to result from these early developmental disruptions has been observed in the majority of children with ASD (10/11) in one post-mortem study, including in the posterior superior temporal cortices where auditory cortex is located (Stoner et al., 2014, also see Hutsler and Casanova, 2015 and McFadden and Minshew, 2013). Finally, distorted tonotopic maps in primary auditory cortex have been found in animal models of autism (Anomal et al., 2015; Nagode et al., 2017).

The auditory system is one of the earliest to mature in the course of brain development. Functional connectivity between thalamus and auditory cortices has been observed in neonates using in vivo fMRI (Alcauter et al., 2014; Ferradal et al., 2019), and interhemispheric functional connectivity between auditory cortices is already established in infants born prematurely and scanned before term-equivalent age (Doria et al., 2010), as well as in healthy fetuses undergoing fMRI in utero (Jakab et al., 2014; Thomason et al., 2013). Early maturation makes the auditory network vulnerable to disruption in utero or as a result of complications during birth (Barkat et al., 2011). For instance, interhemispheric auditory functional connectivity has been shown to be reduced in fetuses prior to preterm birth (Thomason et al., 2017). Rotem-Kohavi et al. (2019) observed overconnectivity within the auditory network in neonates whose mothers took selective serotonin reuptake inhibitors to treat depression during pregnancy, compared to those who did not.

Disruption to auditory cortical development likely results in atypical sound processing, with potentially cascading consequences for language and social development. In ASD, atypical cortical sound processing is evident from studies utilizing electroencephalography (EEG; Ferri et al., 2003; Hudac et al., 2018; Schwartz et al., 2018), magnetoencephalography (MEG; Yoshimura et al., 2016) and fMRI (Gomot et al., 2006; Green et al., 2019, 2017, 2015), and atypical sensitivity to sounds constitutes one of the most frequent sensory symptoms reported, affecting up to 65% of individuals with ASD (Green et al. 2015; Tharpe et al. 2006). Atypical functional connectivity between the thalamus and cortex, in particular auditory cortex, has been observed in multiple fMRI studies of children, adolescents and young adults with ASD, and has been linked to sensory as well as social deficits and repetitive behaviors (Cerliani et al., 2015; Green et al., 2017; Iidaka et al., 2019; Linke et al., 2018; Maximo and Kana, 2019; Mizuno et al., 2006; Nair et al., 2015, 2013; Woodward et al., 2017).

In addition to guiding the early development of cortical topographic organization and relaying sensory information from the periphery to primary sensory cortices, the thalamus and thalamocortical connections play an important role in regulating sleep (Anderson et al., 2005; Cueni et al., 2008; Jan et al., 2009; Steriade et al., 1993). Simultaneous fMRI-EEG studies in adults show substantial changes in functional thalamocortical connectivity during the transition to sleep (Hale et al., 2016; Mitra et al., 2017, 2015; Picchioni et al., 2014; Spoormaker et al., 2010; Tagliazucchi et al., 2012; Tagliazucchi and Laufs, 2014). In ASD, sleep problems are common, reported by 40-80% of individuals (Arazi et al., 2019; Carmassi et al., 2019; Krakowiak et al., 2008; Mannion and Leader, 2014; Reynolds et al., 2019; Richdale and Schreck, 2009; Sivertsen et al., 2012), and are associated with more severe ASD symptomatology and heightened sensory sensitivity (Mazzone et al., 2018; Sikora et al., 2012; Tzischinsky et al., 2018). Together, these findings point toward disrupted thalamocortical connectivity as a possible mechanism underlying both sleep disturbances and sensory sensitivities in ASD. The current study therefore investigated thalamocortical connectivity, sleep problems, and atypical sensory processing relate in a cohort of toddlers and preschoolers with ASD.

We first aimed to examine links between sleep problems and sensory sensitivities in 15 to 65-month-old toddlers and preschoolers with ASD compared to an age-matched typically developing (TD) control group. Next, we assessed whether atypical functional connectivity (estimated from fMRI acquired during natural sleep) between the thalamus and auditory cortices could be observed in this age range, and whether it was related to sleep problems and sensory sensitivities. Given the early development of connections between the thalamus and auditory cortex, we hypothesized that functional connectivity would be increased in preschoolers with ASD and that the strength of thalamocortical functional connectivity would be related to sleep problems and sensory sensitivities. Lastly, we quantified the amplitude of low-frequency fluctuations (ALFF and fALFF, Zou et al., 2008; Zuo et al., 2010) in primary auditory cortex during resting state fMRI. Given that this index provides a measure of BOLD activity, we expected it to be increased in ASD, reflecting amplified sound processing and reduced thalamocortical gating during sleep in the noisy environment of an MRI scanner.

## 2. Results

### 2.1. Sleep problems in preschoolers with ASD are associated with sensory sensitivities

In a cohort of 70 young children with ASD and 46 typically developing (TD) children, ages 15 to 65 months, enrolled in an ongoing longitudinal study of early brain markers of autism (see Methods for inclusion criteria, and Table 1 and Table S1 for demographics), sleep problems were significantly more pronounced in children with ASD compared to TD children (Child Behavior Checklist [CBCL] Sleep Problems T score, *t*(98)=−3.82, *p*<.001, Figure 1A).

**Table 1.** Demographics for full and fMRI cohort (reported with M (mean) ± SD (standard deviation) and [range] when applicable). The ASD and TD groups were matched on age, sex, and in-scanner head motion (RMSD). Information on exact gestational age at birth was missing for 8 children with ASD (2/8 known to be born at term) and 3 TD children; this includes 2 ASD children (one known to be born at term) and 1 TD child in the fMRI cohort.

|  | Full Cohort |  |  | fMRI Cohort |  |  |
| --- | --- | --- | --- | --- | --- | --- |
|  | ASD | TD |  | ASD | TD |  |
| n | 70 | 46 |  | 29 | 30 |  |
| Gestational age at birth (weeks) | 38.4 ± 2.6<br>[31-43] | 39.7 ± 1.1<br>[37-42] | t(103)=3.1<br>p < .05 | 38.5 ± 2.5<br>[31-43] | 39.5 ± 1.2<br>[37-42] | t(54)=1.84<br>p = .07 |
| Age (months) at behavioral assessment | 34.7 ± 12.4<br>[17-64] | 32.6 ± 15.2<br>[15-64] | t(114)=-.82<br>p = .42 | 33.3 ± 9.8<br>[17-54] | 34.1 ± 14.3<br>[15-64] | t(57)=.25<br>p = .81 |
| Age (months) at MRI scan | n/a | n/a | n/a | 34.7 ± 10.1<br>[18-56] | 34.5 ± 14.8<br>[16-65] | t(57)=-.05<br>p = .96 |
| Sex (# female) | 16 | 21 | $\chi^2 = 6.58$<br>p = .01 | 9 | 13 | $\chi^2 = .96$<br>p = .33 |
| ADOS-2 SA | 11.8 ± 4.6<br>[3-20] | n/a | n/a | 12.0 ± 4.9<br>[3-19] | n/a | n/a |
| ADOS-2 RRB | 3.3 ± 2.1<br>[0-8] | n/a | n/a | 3.5 ± 2.2<br>[0-8] | n/a | n/a |
| ADOS-2 Total | 15.0 ± 5.5<br>[4-26] | n/a | n/a | 15.5 ± 5.7<br>[6-26] | n/a | n/a |
| RMSD run1 | n/a | n/a | n/a | .12 ± .04<br>[.047-.188] | .11 ± .04<br>[.046-.172] | t(57)=-1.04<br>p = .30 |
| RMSD run2 | n/a | n/a | n/a | .12 ± .045<br>[.046-.24] | .10 ± .03<br>[.05-.176] | t(57)=-1.18<br>p = .24 |

**Figure 1.**
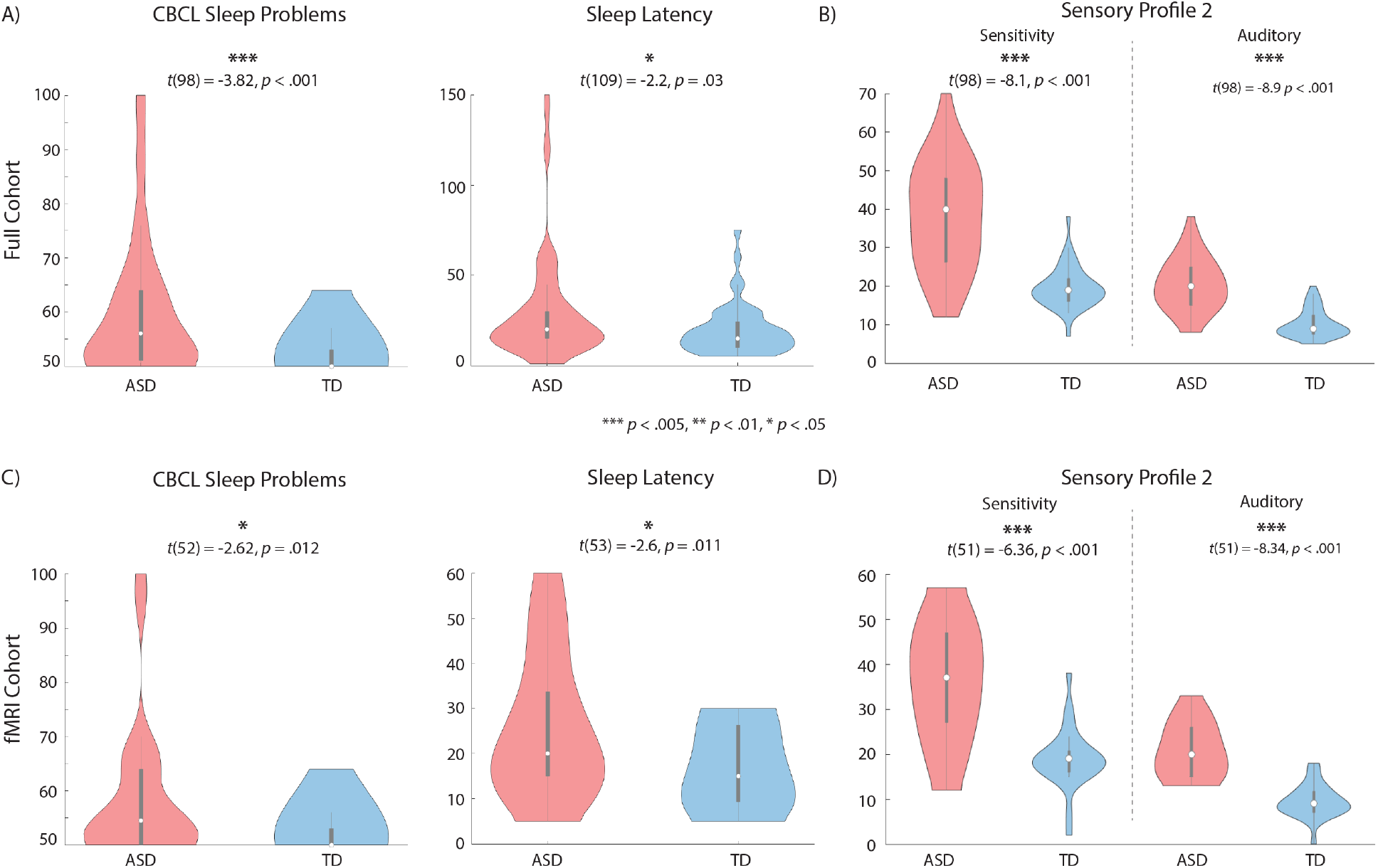
Greater sleep problems and sensory sensitivity, including auditory sensitivity, in toddlers and preschoolers with ASD. **(A, C)** More severe sleep problems (as measured using the CBCL Sleep Problems scale, T scores) and prolonged sleep latency (time, in minutes, it takes a child to fall asleep, as reported on an in-house sleep questionnaire that asked caregivers about their child’s sleep habits in the past few weeks prior to participating in the study) are reported for young children with ASD compared to age and sex matched TD children, in Full Cohort (A) and the subgroup of children with successful fMRI scans (fMRI Cohort, C). **(B, D)** Toddlers and preschoolers with ASD show greater sensory sensitivity and more severe auditory processing symptoms as measured using the Sensory Profile 2 (T scores), in Full Cohort (B) and the subgroup of children with successful fMRI scans (fMRI Cohort, D).

Since the summary CBCL Sleep Problems scale might not be an ideal measure of sleep quality (Gregory et al. 2011), we also tested for group differences in each individual item from the CBCL that assesses sleep (see Table S2). Toddlers and preschoolers with ASD were reported to have more “trouble sleeping” (χ^2^=16.4, *p*<.001) and were more likely to “resist bedtime” (χ^2^=15.3, *p*<.001), to “wake up at night” (χ^2^=11.0, *p*=.004), and to be “overtired” (χ^2^=5.99, *p*=.05) and “sleepless” (χ^2^=12.03, *p*=.001). Additionally, responses from an in-house Sleep Questionnaire (adapted for preschool age from the Brief Infant Sleep Questionnaire (BISQ), Sadeh, 2004) were assessed (Table S2). Toddlers and preschoolers with ASD were reported to take significantly longer to fall asleep (sleep latency/“time to fall asleep”: ASD mean=30.3 minutes (SD=30.1), TD mean=19.7 minutes (SD=14.4), *t*(109)=−2.2, *p*=.03).

As expected, the Sensory Sensitivities score as well as the Auditory Processing score from the Toddler or Child Sensory Profile 2 (Dunn, 2014) were significantly higher in the ASD than the TD group (*p*<.001, Figure 1B). The CBCL Sleep Problems T score positively correlated with the Sensory Profile 2 Sensory Sensitivity quadrant in the ASD group (*r* = ․35, *p* = ․008; controlling for age, Figure 2A and Table S2), with higher scores on each scale corresponding to greater impairment. The correlation with the Sensory Profile 2 Auditory Processing score was not significant (*r* = ․12, *p* = ․37; controlling for age, Table S3).

**Figure 2.**
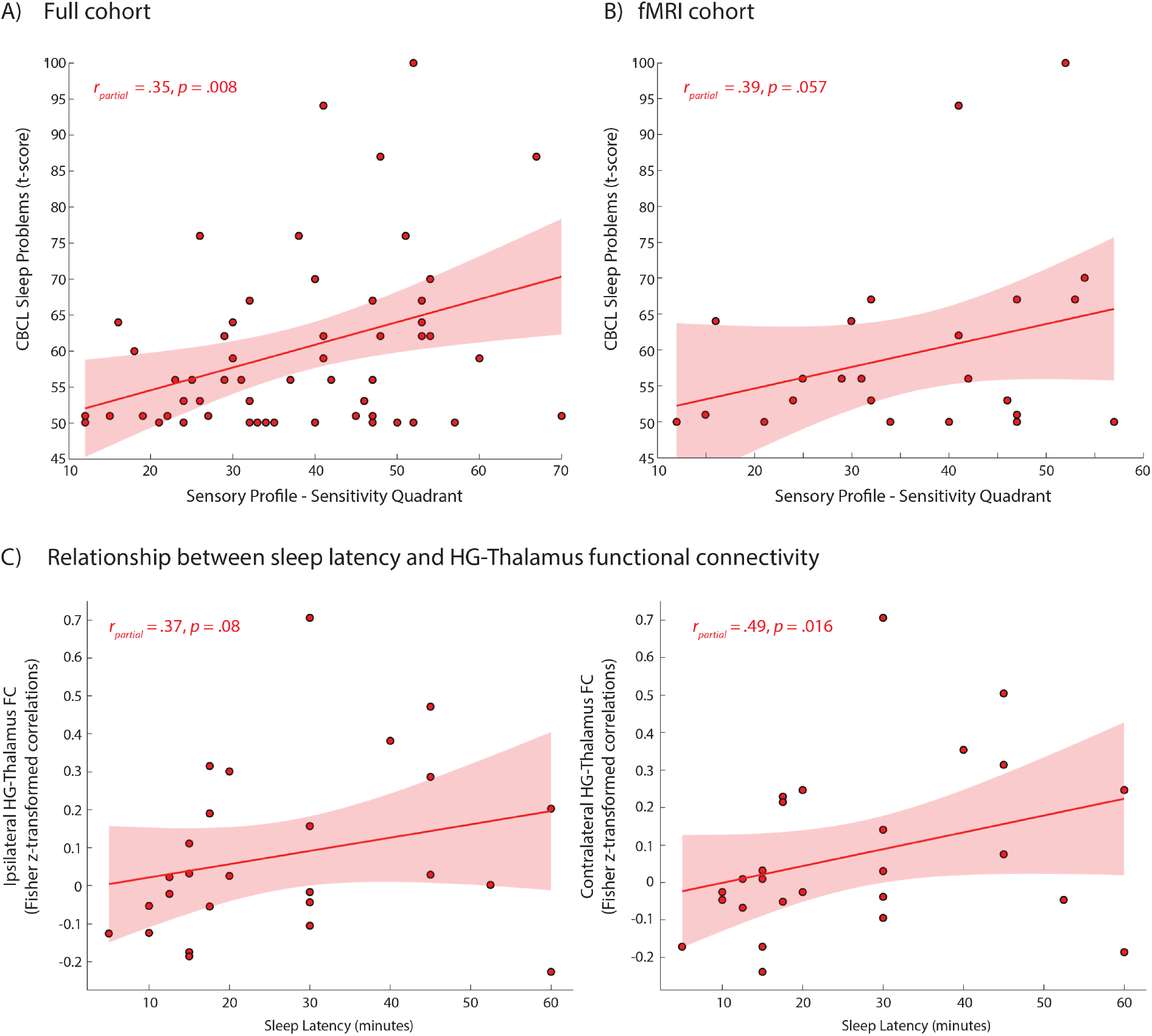
Correlations between sensory sensitivity, sleep problems, and HG-Thalamus FC. **(A, B)** The Sensory Profile Sensitivity Score correlates positively (partial correlation controlling for age) with sleep problems as quantified using the CBCL Sleep Problems T Score in the full (A) and fMRI only (B) ASD group. Correlations were not assessed for the TD group due to the narrow distribution of scores in typically developing children for these measures. **(C)** Sleep latency correlates positively (partial correlations controlling for in-scanner head motion [RMSD] and age) with contralateral FC between the thalamus and HG in the ASD group. Scatterplots show zero-order correlations.

A subsample of children (see 4.1 and Table 1) successfully underwent natural sleep MRI including a high resolution structural T1-weighted image and two multiband echo-planar imaging (EPI) fMRI scans of 6-minute duration each. The ASD and TD groups with fMRI data (henceforth, fMRI cohort) did not differ on age, sex, and head-motion during the fMRI scans, and included 29 ASD and 30 TD participants. Children with successful fMRI scans did not significantly differ on any of the included Sensory Profile or CBCL sleep measures from those without MRI (neither in the ASD nor TD group, Table S4). As in the full cohort, children with ASD and successful fMRI scans had more sleep problems compared to TD peers as measured using the CBCL (Sleep Problems T score: *t*(52)=−2.62, *p*=.012, “trouble sleeping”: χ^2^=9.62, *p*=.008, “resists bedtime”: χ^2^=9.73, *p*=.008, and, marginally, “wakes up at night”: χ^2^=4.95, *p*=.08). Similarly, sleep latency was prolonged in children with ASD (ASD mean=26.9 minutes (SD=16.1), TD mean=17.4 minutes (SD=10.4), *t*(53)=−2.6, *p*=.011, Figure 1C).

Toddlers and preschoolers with ASD in the fMRI cohort also showed greater sensory sensitivities and more severe auditory processing symptoms as assessed using the Sensory Profile 2 (both *p*<.001, Figure 1D, Table S2). Similarly, the correlation between the Sensory Profile 2 Sensory Sensitivity Quadrant score and the CBCL Sleep Problems T score was positive (*r* = ․39, *p* = ․057), consistent with the full cohort data (Figure 2B). The correlation between the Sensory Profile 2 Auditory Processing score and the CBCL Sleep Problems T score was not significant (r=.23, p=.28; partial correlations controlling for age, Table S3).

### 2.2. FC between the thalamus and HG is increased in preschoolers with ASD

Functional connectivity analyses were carried out for left and right Heschl’s Gyrus (HG) and left and right thalamus regions of interests (ROIs). FC between the thalamus and HG was significantly increased between right HG and right and left thalamus in the ASD compared to the TD group (*t*(57)=−2.7, *p*=.01 and *t*(57)=−2.8, *p*=.007, respectively, Figure 3A). FC was also higher between left HG and left and right thalamus in the ASD group but the group difference was not significant (*t*(57)=−1.3, *p*=.19 and *t*(57)=−1.76, *p*=.08, respectively). Ipsilateral and contralateral FC between HG and the thalamus were significantly higher in the ASD group (*t*(57)=−2.5, *p*=.015 and *t*(57)=−2.32, *p*=.024, respectively). In many TD children FC estimates were close to zero or negative in line with previous reports of decreasing thalamocortical FC with anticorrelations frequently observed during deep sleep (Picchioni et al., 2014). These results remained similar when global signal regression (GSR) was included during fMRI data denoising, with FC between left HG and left and right thalamus additionally showing significant group differences after GSR (Figure S1). There was no significant correlation between functional connectivity strength and age in the ASD or TD group, or when carrying out correlations across the combined cohort for any of the auditory-thalamic FC estimates (all *r* < ․2, *p* > ․2).

**Figure 3.**
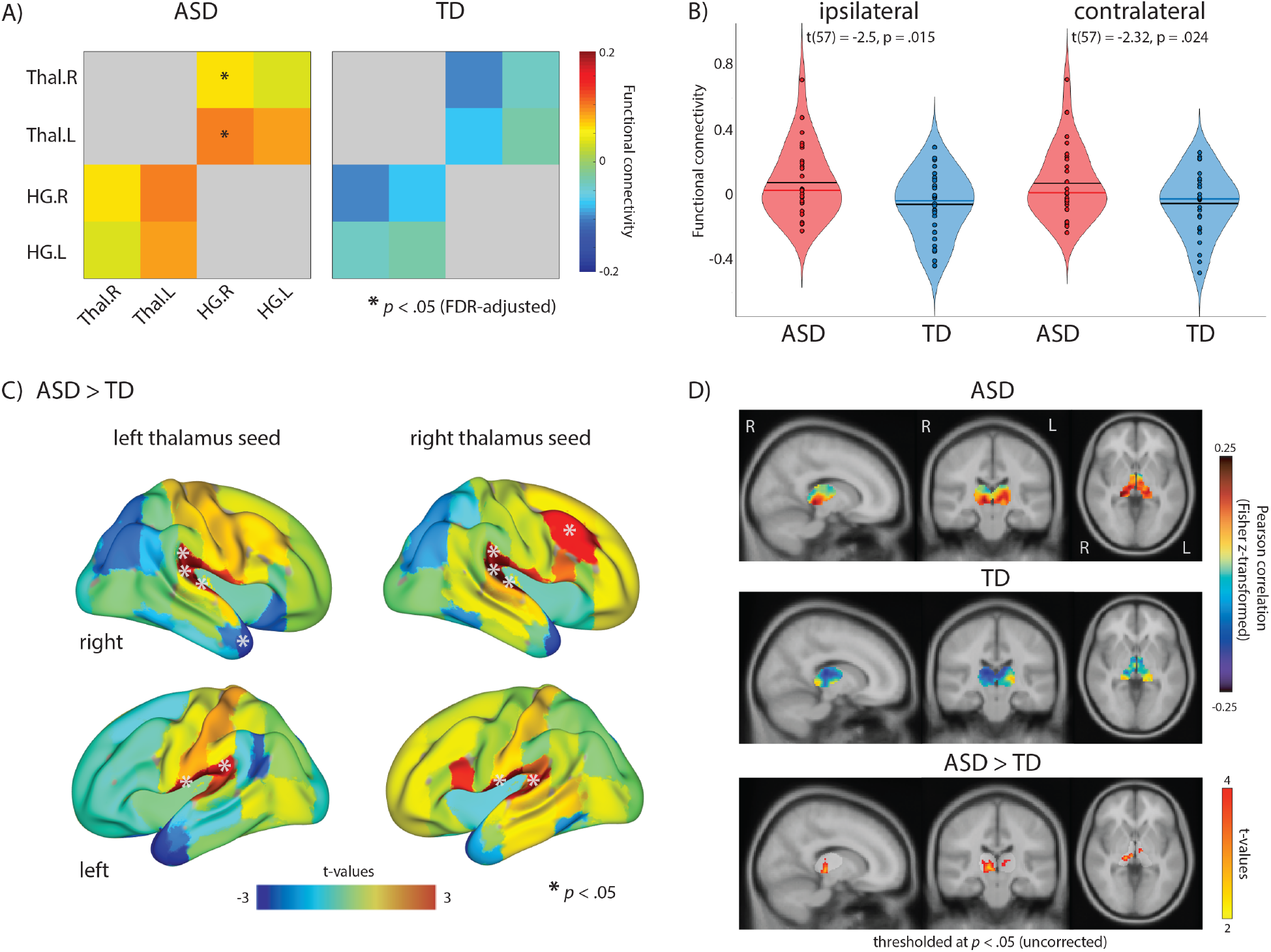
Thalamic-HG overconnectivity in young children with ASD. **A)** Toddlers and preschoolers with ASD show overconnectivity between thalamus and HG during natural sleep fMRI compared to age, motion and sex matched TD children (* in the ASD matrix mark ROI pairs with significant group differences, *p* < ․05, Benjamini-Hochberg FDR-adjusted for multiple comparisons). **B)** Ipsilateral and contralateral FC was similarly elevated in toddlers and preschoolers with ASD and used for behavioral correlations in subsequent analyses. **C)** Seed-to-ROI analyses show that thalamocortical overconnectivity is most pronounced for auditory regions. Left and right thalami were used as seeds and all cortical Harvard-Oxford atlas ROIs as targets. t-values for the ASD>TD comparison are shown with * marking significant (p < ․05, uncorrected) group differences in FC. **D)** Thalamic-HG overconnectivity in the ASD group appears to be driven by the posterior section of the thalamus. Left and right HG were used as seeds and all voxels in the thalamus as targets.

In order to assess whether overconnectivity was specific to auditory regions, seed-to-ROI analysis was carried out using the left and right thalamus as a seed and all cortical Harvard-Oxford ROIs as targets. Based on the pattern of group differences in FC (Figure 3B), results suggest overconnectivity between the thalamus and cortex is most pronounced in the temporal lobe around primary auditory cortex with right HG having the highest effect size among all comparisons for the left thalamus seed (Cohen’s d = ․73). Left HG similarly was among highest effect sizes for both left and right thalamus seeds (Cohen’s d=.46 and d=.34, respectively).

Lastly, we assessed whether overconnectivity between thalamus and primary auditory cortex was driven by specific thalamic sub-regions. Using left and right HG as seeds, Pearson correlations were calculated for every voxel in the thalamus ROI. Results are shown in Figure 3C, with the pattern of FC suggesting that overconnectivity is strongest for the posterior regions of the thalamus, potentially overlapping with the medial geniculate nucleus, and corresponding to the functional parcellations of the thalamus derived from functional thalamocortical connectivity patterns previously, including in infants (Hwang et al., 2017; Kumar et al., 2017; Toulmin et al., 2015).

### 2.3. BOLD signal amplitude is increased in auditory cortex during natural sleep in toddlers and preschoolers with ASD

The amplitude of low frequency fluctuations (ALFF) measures the power of the BOLD signal within a low frequency range and is thought to reflect the amplitude of regional activity (Zuo et al. 2008; Zuo et al. 2010). ALFF (*t*(57)=2.4, *p*=.02,) and fALFF (*t*(57)=1.9, *p*=.06) were higher in HG in the ASD compared to the TD group (Figure S3). Neither ALFF nor fALFF correlated significantly with Sensory Profile Sensory Sensitivity or Auditory Processing Scores or with the CBCL Sleep Problems T-score in children with ASD.

## 3. Discussion

We investigated the relationship between auditory thalamocortical FC, sleep problems, and sensory sensitivities in toddlers and preschoolers with ASD. Elevated sleep problems and sensory sensitivities, as reported by caregivers, were positively correlated in children with ASD, and increased sleep latency was associated with higher thalamocortical connectivity during natural sleep. Our findings support a model of both atypical sensory processing and sleep problems being linked to early neurodevelopmental disturbances of thalamocortical connectivity.

### 3.1. Atypical thalamocortical FC may underlie both sensory sensitivities and sleep problems in young children with ASD

Sensory over-responsivity can interfere with sleep (Cortesi et al., 2010; Reynolds and Malow, 2011), accounting for correlations reported between sleep problems and sensory sensitivities in ASD (Tzischinsky et al., 2018). The mechanisms behind sensory sensitivities and sleep problems however remain unclear. In this study, we focused specifically on investigating FC between the thalamus and auditory cortex and its association with sleep problems and sensory sensitivities for a number of reasons: 1) connections between the thalamus and auditory cortex are established early in development (Alcauter et al., 2014; Ferradal et al., 2019) making them particularly vulnerable to disruption in utero (Barkat et al., 2011); 2) the formation of tonotopic maps in auditory cortex is guided by thalamocortical connections in utero and its disruption can result in atypical sound processing, as has been shown in animal models of ASD (Anomal et al., 2015; Nagode et al., 2017); 3) altered sensitivity to, and atypical cortical processing of sounds is very common in ASD and has been linked to reduced modulation of thalamocortical FC (Green et al., 2017); and 4) a relationship between atypical sensory gating by the thalamus during sleep as reflected by increased thalamocortical connectivity and elevated BOLD amplitude was likely to be most obvious in auditory cortex given sleep fMRI is collected in the presence of substantial noise produced by the MRI scanner.

Similar to findings in older children and adolescents with ASD who underwent resting state fMRI while awake (Linke et al. 2018; Nair et al. 2015; Woodward et al. 2017), auditory-thalamic FC was elevated in toddlers and preschoolers with ASD scanned during natural sleep in the current study. While auditory-thalamic FC was positively associated with sleep problems, particularly the time it takes children to fall asleep, it did not correlate with sensory sensitivities. The Sensory Profile may not have quantified atypical auditory processing in ASD with high reliability (Schulz and Stevenson, 2019), as it relies on caregiver report and some questions targeting modality-specific processing also tap into non-sensory aspects of behavior. Previous studies employing sensory stimulation fMRI designs (e.g. Green et al. 2017) have found thalamocortical overconnectivity in ASD to be associated with sensory over-responsivity (quantified as a composite score derived from the Short Sensory Profile (Dunn, 1999) and Sensory Over-Responsivity (SensOR) Inventory (Schoen et al., 2008)). The lack of a relationship between sensory sensitivities as broadly assessed with the Sensory Profile and auditory thalamocortical FC is thus likely to reflect the Sensory Profile psychometric limitations.

### 3.2. Evidence for early development of atypical auditory-thalamocortical FC

Auditory-thalamic FC did not correlate with age, suggesting that the observed overconnectivity reflects early neurodevelopmental disruptions preceding toddler age. Postmortem histology and studies in autism animal models provide evidence for altered establishment of thalamocortical projections and topographic sensory maps in utero potentially as a result of atypical subplate function (Barkat et al., 2011; Constantin et al., 2020; Hoerder-Suabedissen et al., 2013; Hutsler and Casanova, 2015; Kanold et al., 2019; McFadden and Minshew, 2013; Molnár et al., 2020; Nagode et al., 2017; Serati et al., 2019). Recent findings from an infant sibling fMRI study support this interpretation, showing increased FC between the thalamus and somatosensory cortex in six-week old infants at high familial risk of ASD (Nair et al., preprint: https://www.biorxiv.org/content/10.1101/2020.06.07.139147v1), which was associated with increased ASD symptoms at 36 months. Relatedly, in a large group of at-risk infant siblings, Swanson et al. (2017) reported that thalamus volumes at 12 months differentially predicted language skills at 24 months in those with ASD compared to those with language delay or without familial risk. Abnormalities in auditory cortical processing (Kolesnik et al., 2019) as well as increased prevalence of sleep problems (Humphreys et al., 2014; Nguyen et al., 2018) tied to differences in brain development (MacDuffie et al., 2020) have also been observed in infant sibling studies of ASD, further strengthening the notion that underlying neurodevelopmental disruptions occur very early.

### 3.3. Possible mechanisms: Atypical modulation of thalamocortical FC during awake and sleep states and reduced sensory gating

Unlike in older children and adults scanned awake, average FC between auditory cortex and the thalamus was close to zero or negative in TD toddlers and preschoolers in our study, with overconnectivity in the ASD group driven by a positive shift in correlation magnitudes. Functional connectivity between cortex and subcortical structures changes substantially during sleep, with a reduction in thalamocortical connectivity observed in fMRI studies conducted during deep sleep in adults (Picchioni et al. 2014). Mitra et al. (2017) scanned asleep young children (from 6 to 24 months) and compared the lag pattern of FC to that of adults scanned awake and asleep. During N3 sleep, the BOLD responses of the thalamus and cortex showed increased lag compared to wakefulness, in both sleeping 2-year-olds and sleeping adults. While the authors did not report zero-lag FC of the thalamus, increased lag of thalamic BOLD timeseries during N3 sleep is likely to result in negative or reduced thalamocortical FC compared to wakefulness. Increased FC in the ASD group observed in the current study may therefore reflect a lack of thalamocortical modulation during sleep. It is not possible to discern from the current study whether those children with increased thalamocortical connectivity during deep sleep would also show heightened thalamocortical connectivity while awake. However, findings from previous studies show that thalamocortical overconnectivity in ASD is also present in the awake state. Similarly, in schizophrenia, thalamocortical overconnectivity during awake state has been observed in multiple studies (Chen et al., 2019; Skåtun et al., 2018; Woodward et al., 2012), and is associated with reduced sleep spindle density (Baran et al., 2019). Baran et al. conclude that reduced inhibition by the thalamic reticular nucleus results in both reduced sensory gating (reflected by increased thalamocortical FC) and sleep spindle deficits. Reduced sleep spindle density and duration has also been observed in 2 to 6-year-old children with ASD (Farmer et al., 2018). During wakefulness, children with ASD not only show thalamocortical overconnectivity but also less modulation of thalamocortical connectivity in response to sensory stimulation (Green et al., 2017) and less habituation of auditory cortical processing when presented with aversive auditory stimuli (Green et al., 2019). Additionally, in healthy adults, activity in auditory cortex decreases during sleep (Czisch et al., 2002). In our study, ALFF – a measure of the amplitude of BOLD fluctuations – in HG was significantly higher during sleep in toddlers and preschoolers with ASD, potentially reflecting heightened ongoing sound processing in the auditory cortex or a lack of habituation to the scanner noise. In combination with the previous literature and increased auditory-thalamic FC, these findings support an explanation of reduced sensory gating during sleep in ASD. Atypical modulation of thalamocortical functional connectivity during awake and sleep states might thus offer an explanation for both sensory sensitivities and sleep problems in ASD.

In the absence of simultaneous EEG recordings – which cannot be obtained in sleeping toddlers or preschoolers during fMRI without risking substantial data loss – exact sleep stage is difficult to objectively determine in natural sleep fMRI studies. Note that obtaining MRI data with sufficient data quality in sleeping toddlers and preschoolers is practically impossible unless a child is in deep, motionless sleep. We nevertheless conducted a number of additional analyses suggesting that auditory-thalamic overconnectivity in preschoolers with ASD was not driven by differences in sleep stage (see Supplementary Material and Figure S4). These additional analyses, however, cannot rule out other qualitative differences in sleep. This is supported by a number of EEG studies showing atypical sleep architecture in young children with ASD (e.g. Arazi et al., 2019; Farmer et al., 2018; Lehoux et al., 2019; Page et al., 2020). Particularly, atypical slow wave activity (Arazi et al. 2019; Lehoux et al. 2019) and reduced sleep spindle generation (Farmer et al. 2018) as observed with EEG in young children with ASD has been associated with altered thalamocortical FC during deep sleep (Baran et al. 2019). This is, however, fully consistent with our conclusion of sleep problems in ASD being associated with atypical auditory-thalamic FC and sensory sensitivities. Given the consequences that disrupted sensory processing and sleep might have on development, understanding the relationship between mechanisms underlying sleep problems and the emergence of core ASD symptomatology early in life is of crucial importance. Our findings suggest that early developmental abnormalities of thalamocortical connectivity in ASD are linked to both sleep disturbances and sensory problems, laying out a pathway for mechanistic models and ultimately targeted neurobehavioral interventions.

## 4. Methods

### 4.1. Participants

70 young children with ASD and 46 typically developing (TD) children, ages 15 to 65 months, were enrolled in an ongoing longitudinal study of early brain markers of autism. Children with a (suspected) diagnosis of autism were referred from specialty autism clinics, state-funded early education and developmental evaluation programs, local pediatricians, service providers, and community clinics; TD children were recruited from the community. Participants were excluded for comorbid ASD-related medical or genetic conditions (e.g., epilepsy, fragile X or Rett syndrome, tuberous sclerosis), or other neurological conditions (e.g., cerebral palsy, Tourette syndrome). Participants in the TD group were also excluded for premature birth (< 36 weeks gestational age at birth), and personal or family history (first degree relatives) of ASD, intellectual disability, or other heritable psychiatric or neurological disorders.

All participants were safety-screened for MRI contraindications (e.g., ferrous material in body). Consent was acquired from caregivers, and families were compensated for their time. The research protocol was approved by the institutional review boards of the University of California San Diego and San Diego State University. MRI data were acquired during natural sleep (see SI) from 76/116 children (42 ASD, 34 TD). MRI data were missing in 40 children, as a result of inability to fall or stay asleep in the scanner (29 participants: 21 ASD, 8 TD) or because they did not return for their MRI appointment (11 participants: 7 ASD, 4 TD). 17 MRI datasets were further excluded from analyses due to excessive motion or missing one of the two fMRI runs (n=13 ASD, n=1 TD) or to achieve age and sex matching (n=3 TD) between the ASD and TD groups. The final ASD and TD groups with fMRI data did not differ on age, sex, and head-motion during the fMRI scans, and included 29 ASD and 30 TD participants. Demographics for all participants included in behavioral (full cohort) and fMRI (fMRI cohort) analyses are summarized in Table 1 and Table S1.

### 4.2. Diagnostic and Behavioral Assessments

All participants in the ASD group received a diagnosis of ASD (or clinical best estimate in children younger than age 3 years, Ozonoff et al., 2015) based on the DSM-5 criteria (American Psychiatric Association, 2013), supported by the Autism Diagnostic Observation Schedule, Second Edition (ADOS-2, Lord et al., 2012, Table 1), administered by research-reliable clinicians, the Autism Diagnostic Interview-Revised (ADI-R, for children older than 36 months), and expert clinical judgment. Parents also completed the Social Communication Questionnaire (SCQ, Current form, Lord and Rutter, 2003), a screener for autism spectrum disorders, with no TD participants exceeding the cut-off score of 15 (all TD scores ≤ 10). Measures summarizing sleep problems were obtained from the Preschool Child Behavior Checklist (CBCL) and from an in-house Sleep Questionnaire (adapted for toddlers and preschoolers from the Brief Infant Sleep Questionnaire (BISQ), Sadeh, 2004). The CBCL Sleep Problems T score, the six individual items it is derived from (Gregory et al., 2011) and 7 items from the Sleep Questionnaire were included in analyses (see SI for details on CBCL sleep items). To quantify sensory symptoms and in particular sound sensitivities, the Toddler or Child Sensory Profile 2 (Dunn, 2014) was administered to caregivers, and the “sensitivity” quadrant score and auditory processing score were used in analyses (with greater scores corresponding to greater impairment).

ASD and TD group differences in the CBCL Sleep Problems T score, individual CBCL sleep items (see SI), the Sleep Questionnaire items, and the Sensory Profile scores were tested using independent samples two-tailed t-tests or likelihood ratio chi square tests for the full cohort and separately for the subgroup of children with fMRI data. Given the restricted range of scores in the TD group, Pearson correlations were carried out to test for a relationship between sensory sensitivities (Sensory Profile Sensitivity quadrant and Auditory Processing score) and sleep problems (CBCL Sleep Problems T score) in the ASD group only.

### 4.3. Magnetic Resonance Imaging Data Acquisition

In preparation for the scan night, and to optimize MRI data acquisition, a comprehensive habituation protocol was implemented. After an individualized scan night sleep strategy (e.g., time of arrival, approximating home-like sleeping arrangements, including access to a double MRI bed for co-sleeping families, lighting in the MRI suite, etc.) was developed for each child, based on the typical bedtime routines and habits assessed in advance, families were instructed to practice inserting soft foam child-size earplugs and to play recordings of the MRI sequences employed in the study for at least a week prior to the scan. During the MRI session, children wore earplugs (~30 dB sound attenuation), and MRI-compatible headphones (MR Confon) were used to play white noise at a comfortable listening level to keep the noise level constant and prevent children from waking up during scan transitions. A weighted blanket kept children comfortable and reduced motion, and a caregiver and/or member of the research team remained in the scanner room throughout the session to monitor sleep, motion, and comfort.

Natural sleep MRI data were collected at the University of California San Diego Center for Functional MRI on a GE 3T Discovery MR750 scanner using a Nova Medical 32-channel head coil. A multiband EPI sequence allowing simultaneous acquisition of multiple slices was used to acquire two fMRI datasets (6-minute duration each) with high spatial and temporal resolution (TR=800ms, TE=35ms, flip angle 52°, 72 slices, multiband acceleration factor 8, 2 mm isotropic voxel size, 104×104 matrix size, FOV 20.8cm, 400 volumes per run). Two separate 20s spin-echo EPI sequences with opposing phase encoding directions were also acquired using the same matrix size, FOV and prescription to correct for susceptibility-induced distortions. After completion of the fMRI scans, a fast 3D spoiled gradient recalled (FSPGR) T1-weighted sequence was used to acquire high-resolution structural images (0.8 mm isotropic voxel size, NEX=1, TE/TI=min full/1060ms, flip angle 8°, FOV=25.6cm, matrix=320+320, receiver bandwidth 31.25hz). Motion during structural acquisitions was corrected in real-time using three navigator scans (PROMO, real-time prospective motion correction White et al., 2010), and images were bias corrected using the GE PURE option.

### 4.4. Imaging data preprocessing and denoising

MRI data were preprocessed, denoised and analyzed in Matlab 2015b (Mathworks Inc., Natick, MA, USA) using SPM12 (Wellcome Trust Centre for Neuroimaging, University College London, UK), and the CONN toolbox v17f (Whitfield-Gabrieli & Nieto-Castanon, 2012).

The structural images were converted from dicom to nifti format and were coregistered to the mean functional images, segmented and normalized to MNI space using non-linear registration and the default tissue probability maps included with SPM12. The white matter (WM) probability maps obtained from segmentation of the structural image for each individual subject were thresholded at 0.95 and eroded by 1 voxel. WM and CSF time courses were extracted from the thresholded and eroded masks using aCompCor (Behzadi et al., 2007) for subsequent nuisance regression (see below).

Functional images were corrected for susceptibility-induced distortions using the two spin-echo EPI acquisitions with opposite phase encoding directions and FSL’s TOPUP tools (Smith et al., 2004). Subsequently, functional images were motion-corrected using rigid-body realignment as implemented in SPM12. The Artifact Detection Toolbox (ART, as installed with CONN v17f) was used to identify outliers in the functional image time series from the resulting 6 motion parameters (3 translational and 3 rotational) that had frame-wise displacement (FD) >0.5 mm and/or changes in signal intensity that were greater than three standard deviations. As oscillations due to respiration are prominent in motion parameters derived from multiband EPI realignment (Fair et al. 2018) and would result in unnecessary censoring of large chunks of data in some participants, the thresholds to detect outliers were more lenient than those used for standard resting state fMRI acquisitions with slower TRs. In order to ensure that none of our findings were due to differences in apparent motion between groups, groups were matched on RMSD calculated from rigid-body realignment of the raw data prior to TOPUP correction (Table 1).

Functional images were directly normalized to MNI space with the same non-linear registration as used for the structural images. The MNI template rather than an age-appropriate template was used for normalization due to the wide age range and cross-sectional analyses conducted. Data were visually inspected for quality by two members of the research team (authors AL and BC) independently, at each pre-processing step, including determining successful normalization. Voxel timeseries were converted to percent-signal change separately for each EPI acquisition. Since all analyses were run on averaged voxel time series within pre-defined ROIs, no prior smoothing was applied to the data. Band-pass filtering using a temporal filter of 0.008 to 0.08 Hz was carried out as part of the nuisance regression (“simult” option in the CONN toolbox) which also included scrubbing of the motion outliers detected by the ART toolbox, and regression of the 6 motion parameters and their derivatives, as well as the first five PCA component time series derived from the CSF and white matter masks. The residuals of the nuisance regression were then used for all subsequent analyses. Analyses were repeated including the global signal, calculated as the average timeseries of all voxels in the brain, as a nuisance regressor (see Figure S1).

### 4.5. Functional MRI Analyses

#### 4.5.1. Regions of interest

Analyses were carried out within left and right Heschl’s Gyrus (HG) and left and right thalamus regions of interests (ROIs) extracted from the Harvard-Oxford atlas provided by FSL and the CONN toolbox. Additional functional connectivity analyses included all cortical Harvard-Oxford atlas ROIs to determine whether atypical thalamocortical connectivity was specific to auditory cortices (see below).

#### 4.5.2. Functional connectivity analysis

BOLD time series were concatenated across the two EPI acquisitions and averaged across all voxels within each ROI. First, functional connectivity (FC) between bilateral thalamus and HG was estimated using bivariate Pearson correlation standardized with a Fisher z-transform. Two-tailed independent samples t-tests were used to assess differences in correlation magnitude between pairs of ROIs in the ASD compared to the TD group. Separate tests were also carried out for ipsilateral (average of intrahemispheric thalamus-HG FC) and contralateral (average of interhemispheric thalamus-HG) FC. Functional connectivity (Fisher z) was Pearson correlated with age, separately in each group, to test for any age-related changes in auditory thalamocortical connectivity. We hypothesized that auditory-thalamic FC would be elevated in toddlers and preschoolers with ASD. Results are reported at a threshold of p<.05 (Benjamini-Hochberg FDR adjusted for multiple comparisons). To assess whether atypical thalamocortical connectivity was specific to sensory regions of the brain, a post-hoc analysis also assessed whole brain thalamocortical connectivity using the left and right thalamus as seeds and all Harvard-Oxford cortical ROIs as targets. Independent samples t-tests of the Fisher-z-transformed estimates of FC assessed differences between the ASD and TD groups. Lastly, we assessed in an exploratory analysis whether atypical functional connectivity between HG and the thalamus was driven more strongly by specific regions within the thalamus, involved in auditory processing (Hwang et al., 2017; Kumar et al., 2017; Toulmin et al., 2015). BOLD timeseries within left and right HG were Pearson correlated with the timeseries of every voxel within the thalamus ROIs, and t-tests carried out for the Fisher-z-transformed correlation coefficient of each thalamic voxel to test for ASD-TD differences. To rule out that children in the ASD and TD group might have been scanned during different sleep stages, which could confound functional connectivity estimates, a number of additional analyses were conducted and are described in the Supplementary Information.

#### 4.5.3. Fractional amplitude of low frequency fluctuations

The amplitude of low frequency fluctuations (ALFF) measures the power of the BOLD signal within a low frequency range and is thought to reflect the amplitude of regional neural activity. ALFF was calculated as:

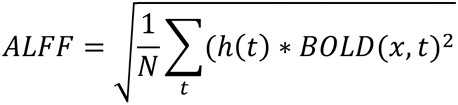

as implemented in the CONN toolbox, with N = the number of volumes, BOLD(x,t) = original BOLD timeseries before bandpass filtering, and h(t) = bandpass filter (see https://web.conn-toolbox.org/measures/other for more detail). The fractional amplitude of low frequency fluctuations (fALFF) was developed to better protect against noise and is a measure of the relative contribution of low frequency fluctuations to the entire frequency range detectable by BOLD-optimized EPI (Zou et al., 2008; Zuo et al., 2010). It was calculated as the power within the low frequency range (0.01 – 0.1 Hz) divided by the total power of the entire frequency spectrum, again using the implementation included with the CONN toolbox:

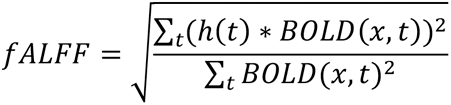

ALFF and fALFF were calculated for each voxel in the brain and extracted and averaged for the left and right HG ROIs. We hypothesized that ALFF and fALFF would be increased in HG in the ASD group, which was tested using independent samples two-tailed t-tests.

### 4.6. Relationship between FC, sleep problems and sensory sensitivities

Partial correlations (correcting for average head motion [RMSD] and age) tested for a relationship between HG-thalamus FC and sensory sensitivities (Sensory Profile Sensitivity Quadrant and Auditory Processing score), the CBCL Sleep Problems T score, and those Sleep Questionnaire items that showed significant ASD-TD group differences in the fMRI cohort. To reduce the number of multiple comparisons, correlations were only conducted for contralateral and ipsilateral FC. Due to the narrow distribution of CBCL and Sensory Profile scores in the TD group, correlations where only assessed for children with ASD. We hypothesized that elevated FC between the thalamus and HG would be related to greater sensory sensitivity and more severe sleep problems in the ASD group. Given relatively small sample size for detecting robust brain-behavior relationships, these results are presented as preliminary and uncorrected for multiple comparisons and need to be interpreted with caution.

## Data Availability

The data that support the findings presented in this manuscript will be available in the NIMH Data Archive (NDA), an NIH-funded data repository (https://nda.nih.gov/). Software used for all analyses are available to researchers for replication.

## Acknowledgments

We thank Lisa Mash, M.S. (San Diego State University) and Tiffany Wang, M.S. (University of California, San Diego) for invaluable assistance with data collection. Our strongest gratitude goes to the children and families who so generously dedicated their time and effort to this research.

