## Supplementary Material for "Sleep problems in preschoolers with autism spectrum disorders are associated with sensory sensitivities and thalamocortical overconnectivity"

#### Supplementary Information

### 1. Supplementary Methods

##### Diagnostic and Behavioral Assessments

*Preschool Child Behavior Checklist (CBCL):* The CBCL is a standardized caregiver questionnaire that aims to identify emotional and behavioral problems in children ages 18 months to 5 years. Six items assess sleep behaviors ("overtired", "sleeps less than most kids", "resists bedtime", "trouble sleeping", "nightmares", "wakes up at night") from which a Sleep Problems T score is calculated. The CBCL Sleep Problems T score only has low correspondence with sleep quality as measured with sleep diaries, actigraphy and polysomnography (Gregory et al., 2011), with some of the individual items (e.g. "trouble sleeping") shown to be more useful for quantifying sleep problems. We therefore also assessed group differences on all individual CBCL sleep behavior items. The CBCL Sleep Problems T score was available for n=100 (59 ASD, 41 TD), and for n=54 (26 ASD, 28 TD) of children who also successfully completed the fMRI scan.

*Sleep Questionnaire:* A sleep questionnaire designed in-house was administered to parents in order to plan the natural sleep MRI session. Items from the in-house Sleep Questionnaire (adapted from the Brief Infant Sleep Questionnaire [BISQ]) that we hypothesized might be related to sensory sensitivities, or that had previously been associated with ASD (e.g. Souders et al. 2009; Reynolds et al. 2019) were extracted. These items included: typical sleep latency ("time to fall asleep" in minutes, recoded to the average if the response was a range), "number of night awakenings" (recoded to the average if the response included a range), "regular bedtime" (yes/no), "resists sleep" (yes/no), "wakes with noise" (yes/no/sometimes), "can be moved from car without waking" (yes/no), "can change clothes while asleep" (yes/no). The Sleep Questionnaire was available for n=111 (67 ASD, 44 TD) and for n=55 (26 ASD, 29 TD), for the full and fMRI cohort, respectively.

*Sensory Profile 2:* The Sensory Profile 2 is a standardized caregiver questionnaire that assesses sensory processing patterns across different modalities, including auditory, visual, tactile and oral processing. Scores were available for n=100 (59 ASD, 41 TD), and for n=53 (25 ASD, 28 TD) of children who successfully completed the fMRI scan.

**Sleep Stage Control Analyses**

MRI data were acquired during natural sleep, with the fMRI scans always acquired first. For a scan to be successful, without a toddler waking up with scanner onset or moving excessively during the scan (e.g. to turn to sleeping on the stomach rather than back – the commonly preferred sleeping position in the studied age range; Sadeh et al., 2009), deep sleep (~N3) needs to be achieved. As such, it is unlikely that sleep stage varied substantially between toddlers, and between the ASD and TD groups. However, without recordings of simultaneous EEG (which is difficult if not impossible to collect in toddlers during natural sleep MRI), sleep stage cannot be unambiguously determined. For this reason, we conducted a number of control analyses to rule out the possibility that our findings were driven by differences in sleep stage during the fMRI scans. First, fMRI scan start time in relation to family arrival time and time the child fell asleep was compared between the ASD and TD groups using independent samples two-tailed t-tests. Secondly, FC between the thalamus and HG was compared between the two subsequent fMRI acquisitions using paired samples t-tests, and an independent-samples two-tailed t-test tested for group differences in the change in FC between the two acquisitions. Lastly, global signal amplitude (the standard deviation of the global signal timeseries) has been shown to reflect levels of vigilance and wakefulness, with a higher amplitude reflecting lower levels of wakefulness (Liu et al., 2017; Wong et al., 2013; Chi Wah Wong et al., 2016). Global signal amplitude in the ASD and TD group was compared using independent samples two-tailed t-tests for both fMRI acquisitions separately.

*Results*

First, there was no significant difference in arrival state (awake/asleep) at the night of the scan between the ASD (awake/asleep = 15/12) and TD (awake/asleep = 20/10) groups ($\chi^2$=.74, $p$=.39). There were also no significant differences in time between arrival at the MRI center and the start of the MRI scan (ASD: 87.2±42.6 minutes; TD: 80.2±55.5 minutes; $t(50)$=-.51, $p$=.61, see Figure S4A), and time between arrival and MRI start time did not correlate with FC between the thalamus and HG for any ROI pair (all $r$<.2, $p$>.2). Similarly, there was no significant difference in time passed between a child falling asleep and the start of the MRI scan between groups (ASD: 23.7±12.8 minutes; TD: 23.2±11.0 minutes; $t(47)$=-.16, $p$=.87, see Figure S4B). Time between falling asleep and the start of the MRI scan further did not correlate with FC between any ROI pair (all $r<.06$, $p>.7$). Second, FC between thalamus and HG did not differ significantly between run1 and run2 for any ROI pair in the ASD or TD group (paired samples t-tests, all $p>.2$), as would have been expected if toddlers with ASD were transitioning through different sleep stages than TD children over the course of the 15 minutes that it took to acquire the two natural sleep fMRI acquisitions. Lastly, global signal amplitude has been shown to correspond closely to vigilance as measured using EEG with higher global signal amplitude reflecting lower levels of vigilance or deeper levels of sleep (Wong et al., 2016; Wong et al., 2013). There were no significant differences in global signal amplitude between the ASD and TD group ($t(57)=-.92$, $p=.36$).

#### 2. Supplementary Tables

**Table S1.** Socioeconomic and sociodemographic characteristics of toddlers and preschoolers included in analyses. Data were missing for three families (2 ASD, 1 TD).

| | ASD n = 68 | | TD n = 45 | | |
|---|---|---|---|---|---|
| | % / Mean | range | % / Mean | range | p-value |
| **% Mothers with College Degree*** | 44% | - | 80% | - | 0.001 |
| **% Fathers with College Degree*** | 53% | - | 75% | - | 0.03 |
| **Income-to-Needs Ratio (mean ± SD)**** | 3.38±2.4 | 0.23-9.37 | 4.20±2.4 | 0.34-9.37 | 0.08 |
| **Median Income by Postal Code, (mean ± SD)***** | $53,094± $18,316 | $26,000- $105,000 | $53,712± $14,980 | $23,584- $92,759 | 0.86 |
| **% Hispanic** | 49% | - | 24% | - | 0.01 |
| **% White** | 59% | - | 67% | - | 0.35 |
| **% Black** | 1.4% | - | 6.5% | - | 0.15 |
| **% Asian** | 4.3% | - | 6.5% | - | 0.62 |
| **% Mixed Race** | 25% | - | 15% | - | 0.2 |

*Maternal and Paternal Educational Level ranged from incomplete high-school education to advanced professional or doctoral degree.
** Income-to-needs ratio is derived by dividing reported household income by the federal poverty level, adjusted for household size. An income-to-needs ratio of 1 indicates living at the federal poverty level.
***Data derived from 2010 census (updated 2018) were used to derive median income by postal code.

**Table S2**. CBCL and SQ sleep items listed for the ASD and TD group. Statistics testing for group differences are reported for the full and fMRI cohort separately. For the CBCL sleep CBCL items: 0=not true, 1=somewhat or sometimes true, 2=very true or often true. For Sleep Questionnaire: smt=sometimes.

| | | **Full Cohort** | | | **fMRI Cohort** | | |
|---|---|---|---|---|---|---|---|
| **Measure** | **Item** | **ASD** | **TD** | | **ASD** | **TD** | |
| CBCL (sleep items) | trouble sleeping | 0 = 27<br>1 = 18<br>2 = 14 | 0 = 26<br>1 = 15<br>2 = 1 | $\chi^2$ = 16.4<br>***p* < .001** | 0 = 12<br>1 = 8<br>2 = 6 | 0 = 16<br>1 = 12<br>2 = 0 | $\chi^2$ = 9.62<br>***p* = .008** |
| | nightmares | 0 = 43<br>1 = 10<br>2 = 6 | 0 = 37<br>1 = 5<br>2 = 1 | $\chi^2$ = 3.59<br>*p* = .17 | 0 = 20<br>1 = 5<br>2 = 1 | 0 = 24<br>1 = 4<br>2 = 0 | $\chi^2$ = 1.79<br>*p* = .41 |
| | overtired | 0 = 43<br>1 = 11<br>2 = 5 | 0 = 35<br>1 = 6<br>2 = 0 | $\chi^2$ = 5.99<br>***p* = .05** | 0 = 19<br>1 = 5<br>2 = 2 | 0 = 22<br>1 = 6<br>2 = 0 | $\chi^2$ = 3.0<br>*p* = .22 |
| | resists bedtime | 0 = 24<br>1 = 19<br>2 = 16 | 0 = 29<br>1 = 11<br>2 = 1 | $\chi^2$ = 15.34<br>***p* < .001** | 0 = 11<br>1 = 6<br>2 = 9 | 0 = 19<br>1 = 8<br>2 = 1 | $\chi^2$ = 9.73<br>***p* = .008** |
| | sleepless | 0 = 38<br>1 = 12<br>2 = 9 | 0 = 38<br>1 = 2<br>2 = 1 | $\chi^2$ = 12.03<br>***p* = .002** | 0 = 19<br>1 = 3<br>2 = 4 | 0 = 25<br>1 = 2<br>2 = 1 | $\chi^2$ = 2.88<br>*p* = .24 |
| | wakes up | 0 = 23<br>1 = 14<br>2 = 12 | 0 = 34<br>1 = 6<br>2 = 1 | $\chi^2$ = 11.02<br>***p* = .004** | 0 = 16<br>1 = 4<br>2 = 6 | 0 = 22<br>1 = 5<br>2 = 1 | $\chi^2$ = 4.95<br>*p* = .08 |
| Sleep Questionnaire | sleep latency (min.) | M = 30.3<br>SD = 30.1 | M = 19.7<br>SD = 14.4 | *t*(109)=-2.2<br>***p* = .03** | M = 26.9<br>SD = 16.1 | M = 17.4<br>SD = 10.4 | *t*(53)=-2.6<br>***p* = .011** |
| | night wakenings (#) | M = .96<br>SD = 1.0 | M = .79<br>SD = .87 | *t*(93)=-.89<br>*p* = .38 | M = 1.0<br>SD = 1.1 | M = .89<br>SD = .96 | *t*(48)=-.38<br>*p* = .71 |
| | regular bedtime | no = 25<br>yes = 44 | no = 12<br>yes = 34 | $\chi^2$ = 1.32<br>*p* = .25 | no = 10<br>yes = 18 | no = 10<br>yes = 20 | $\chi^2$ = .04<br>*p* = .85 |
| | resists sleep | no = 19<br>smt = 37<br>yes = 13 | no = 18<br>smt = 25<br>yes = 3 | $\chi^2$ =4.47<br>*p* = .107 | no = 9<br>smt = 15<br>yes = 4 | no = 12<br>smt = 16<br>yes = 2 | $\chi^2$ = 1.07<br>*p* = .59 |
| | wakes with noise | no = 41<br>smt = 21<br>yes = 6 | no = 29<br>smt = 12<br>yes = 5 | $\chi^2$ = .37<br>*p* = .83 | no = 18<br>smt = 8<br>yes = 1 | no = 17<br>smt = 11<br>yes = 2 | $\chi^2$ = .69<br>*p* = .71 |
| | transfer from car asleep | no = 29<br>yes = 38 | no = 16<br>yes = 28 | $\chi^2$ = .53<br>*p* = .47 | no = 7<br>yes = 20 | no = 10<br>yes = 19 | $\chi^2$ = .49<br>*p* = .49 |
| | change clothes asleep | no = 32<br>yes = 35 | no = 25<br>yes = 19 | $\chi^2$ = .87<br>*p* = .35 | no = 13<br>yes = 14 | no = 16<br>yes = 12 | $\chi^2$ = .48<br>*p* = .50 |

**Table S3**. CBCL Sleep Problems T score and Sensory Profile correlations in the ASD group (n=56 with data from both measures in the full cohort, and n=23 in the fMRI cohort; partial correlations controlling for age).

| Sensory Profile item | Full Cohort | fMRI Cohort |
|---|---|---|
| Sensitivity Quandrant | $r$ = .35, ***p* = .008** | $r$ = .39, $p$ = .057 |
| Auditory Processing | $r$ = .12, $p$ = .37 | $r$ = .23, $p$ = .28 |

**Table S4**. No differences in CBCL Sleep Problems T score, sleep latency or Sensory Profile scores for toddlers with successful fMRI scans compared to those without in the ASD or TD group.

| Measure | ASD | TD |
|---|---|---|
| CBCL Sleep Problems (T score) | $t(58)=.38$, $p=.70$ | $t(39)=-.70$, $p=.49$ |
| Sleep Questionnaire Sleep Latency | $t(77)=.41$, $p=.61$ | $t(42)=1.45$, $p=.15$ |
| Sensory Profile 2 Sensitivity Quadrant | $t(58)=.73$, $p=.47$ | $t(39)=.34$, $p=.74$ |
| Sensory Profile 2 Auditory Processing | $t(58)=-.22$, $p=.82$ | $t(39)=.67$, $p=.51$ |

**Table S5**. Correlations of CBCL Sleep Problems T score, Sleep Questionnaire Sleep Latency and Sensory Profile items with functional connectivity between HG and Thalamus ROIs in the ASD group. Partial correlations (controlling for head motion [RMSD] and age) are reported.

| Measure | ipsilateral | contralateral |
|---|---|---|
| CBCL Sleep Problems (T-score) | *r* = .14, *p* = .49 | *r* = .18, *p* = .38 |
| Sleep Questionnaire Sleep Latency | *r* = .37, *p* = .08 | *r* = .49, ***p* = .016** |
| Sensory Profile 2 Sensitivity Quadrant | *r* = .02, *p* = .94 | *r* = -.02, *p* = .92 |
| Sensory Profile 2 Auditory Processing | *r* = -.06, *p* = .80 | *r* = -.05, *p* = .81 |

#### 3. Supplementary Figures

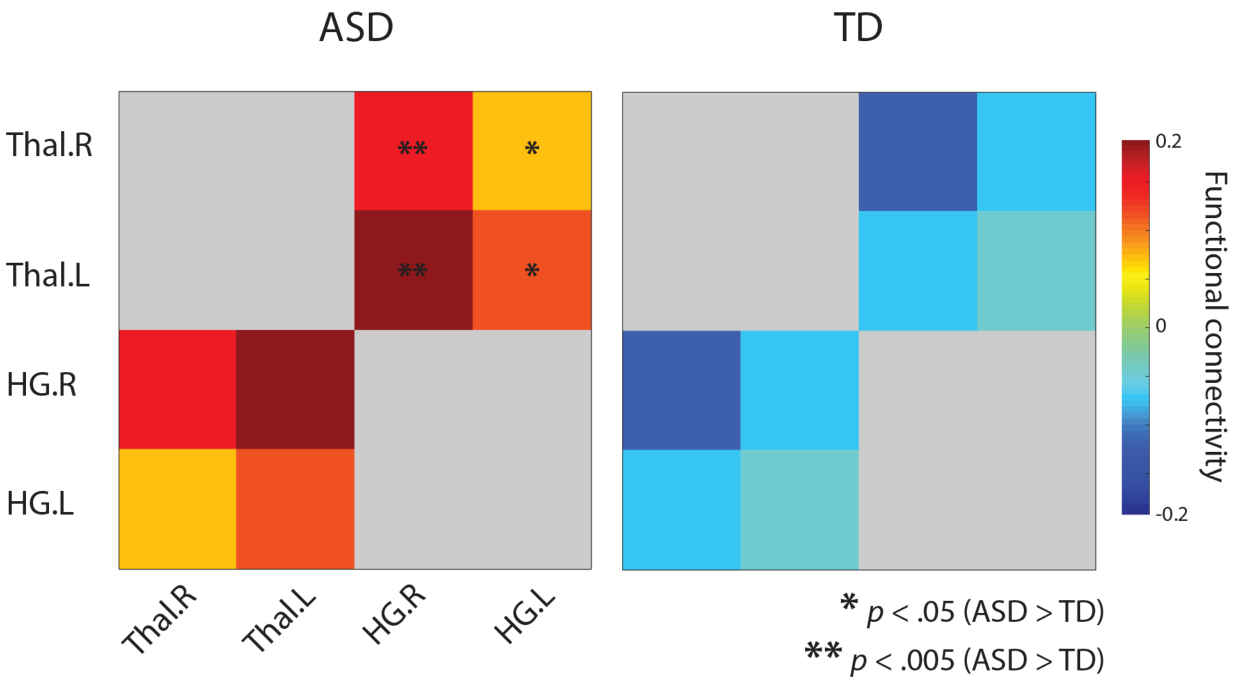


**Figure S1**. FC group differences remain in analyses that include the global signal (mean BOLD timeseries across all voxels) as a nuisance regressor during denoising.

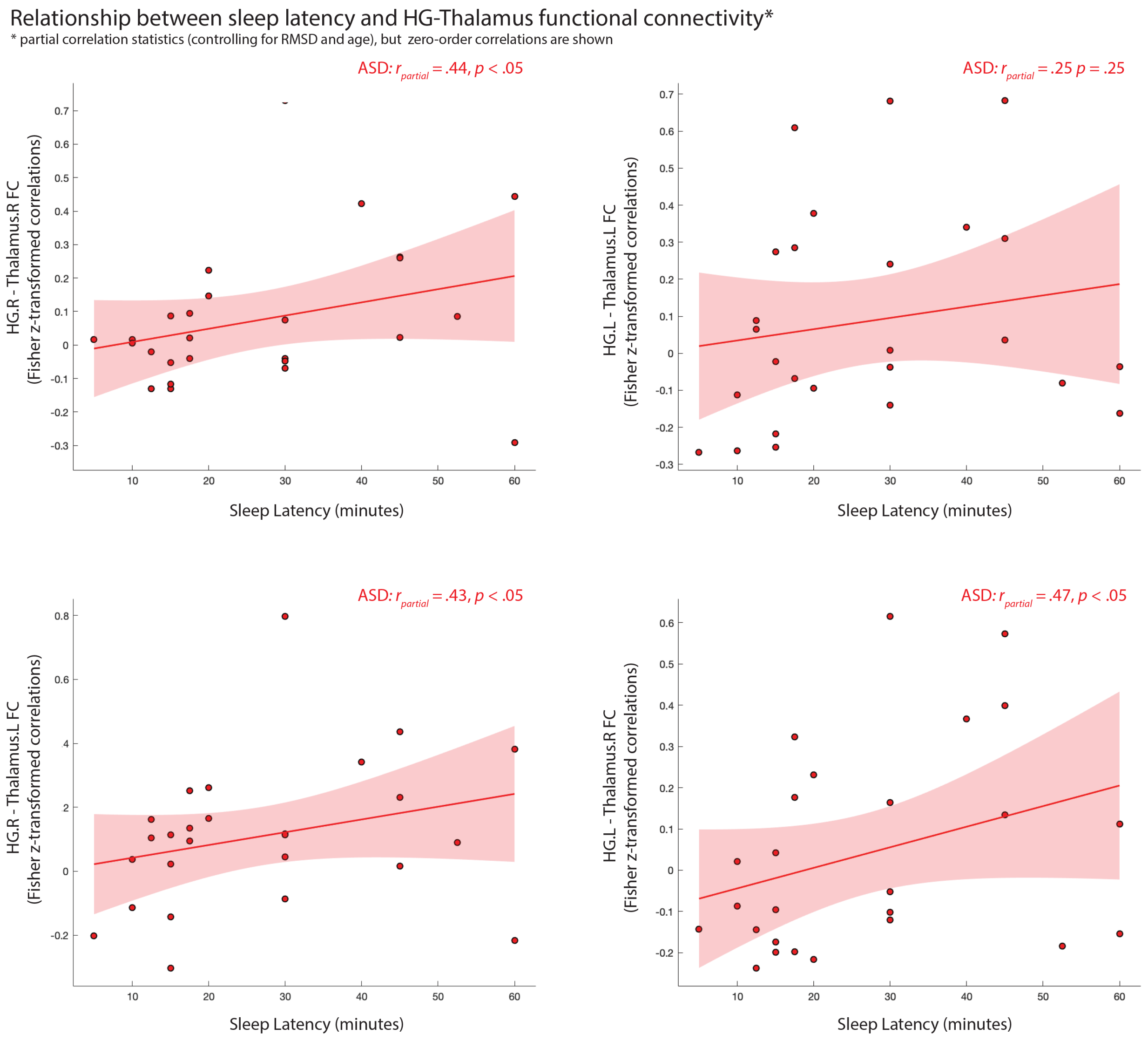


**Figure S2**. Sleep Latency and HG-Thalamus FC correlations for all four ROI pairs separately.

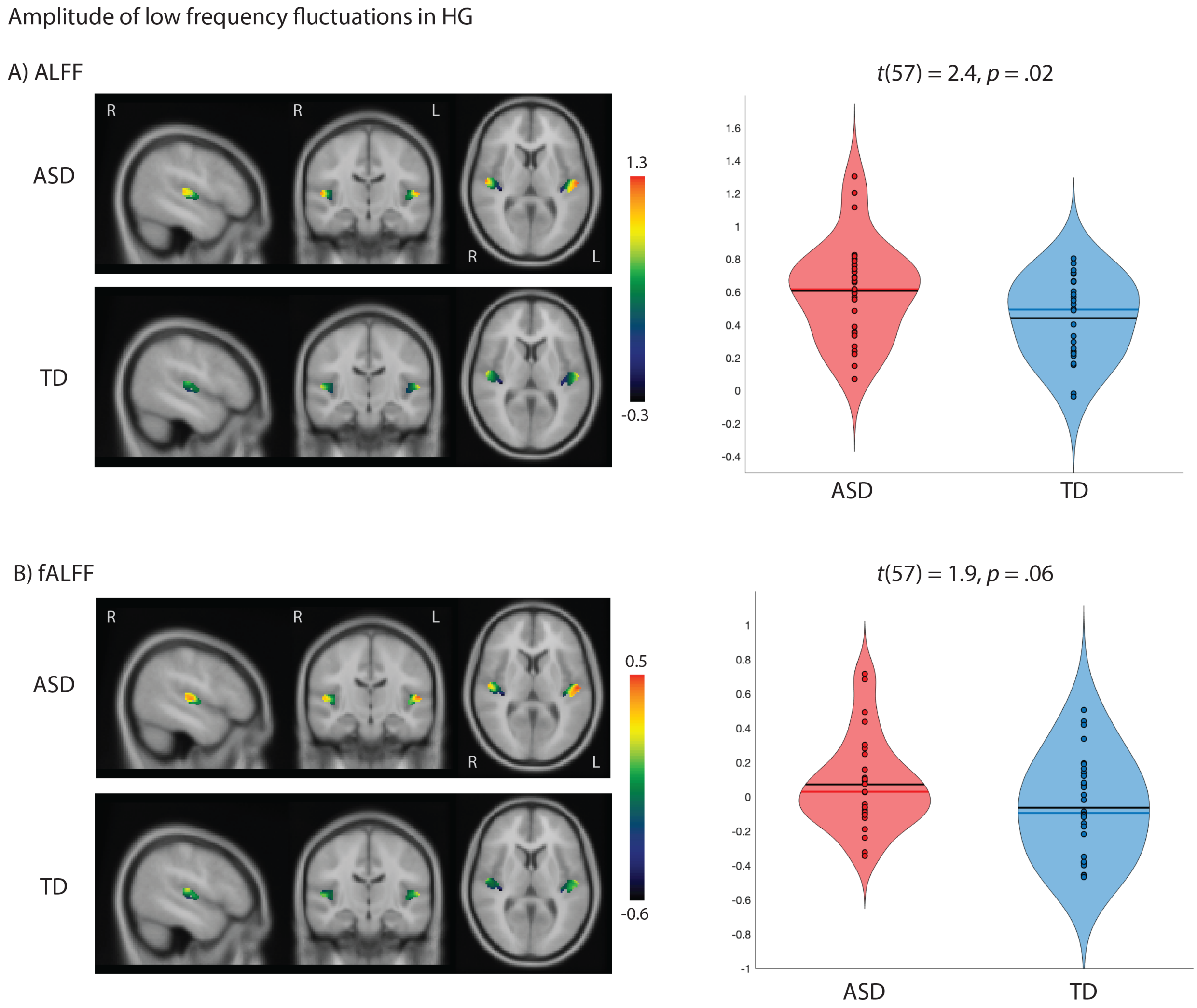


**Figure S3.** The fractional amplitude of low frequency fluctuations (ALFF) in HG (average across hemispheres) is significantly higher in the ASD group compared to the age- and motion-matched TD group.

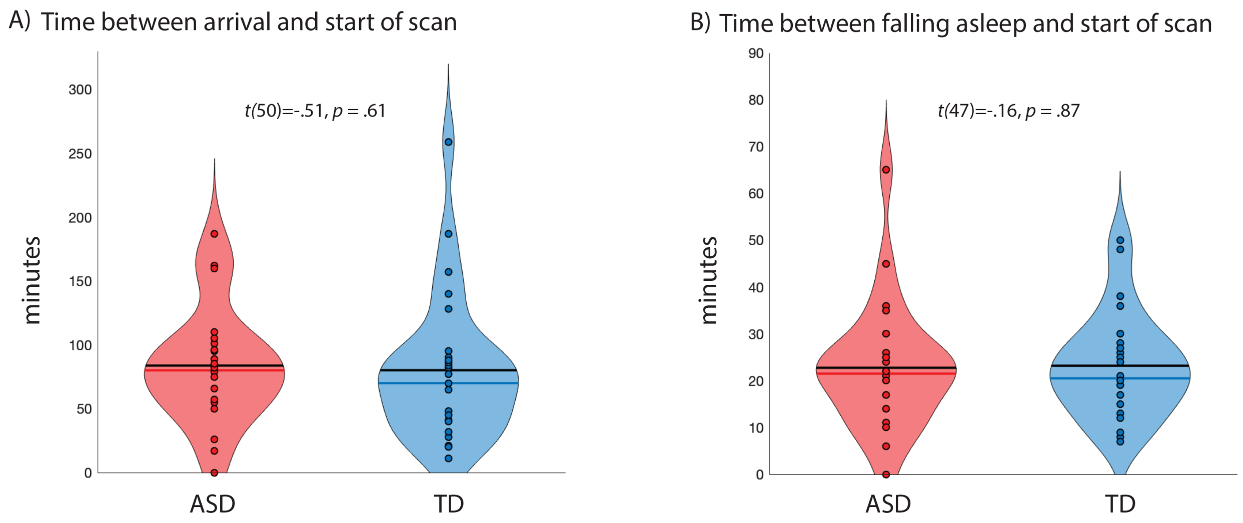


**Figure S4**. There were no significant differences between the ASD and TD groups with successful fMRI scans for the time between arrival at the MRI center and the start of the MRI scan (A), and the time between falling asleep and the start of the MRI scan (B).
